# Detection of urinary metabolites of metabolic pathway disorders by using VTGE and LC-HRMS techniques

**DOI:** 10.1101/814970

**Authors:** Ajay Kumar, Jainish Kothari, Devyani Bhatkar, Manmohan Mitruka, Roshni Pal, Sachin C. Sarode, Nilesh Kumar Sharma

**Affiliations:** Cancer and Translational Research Lab, Dr. D. Y. Patil Biotechnology & Bioinformatics Institute, Dr. D.Y. Patil Vidyapeeth, Pune, Maharashtra, India, 411033; Department of Oral Pathology and Microbiology, Dr. D.Y.Patil Dental College and Hospital, Dr. D. Y. Patil Vidyapeeth, Pimpri, Pune 411018, Maharashtra, India

**Keywords:** Metabolites, Urine, Biomarkers, Metabolic disorders, Mass spectrometry

## Abstract

**Background:** In recent, various human health disorders including cancer, diabetes, neurodegenerative and metabolic diseases are noticed among human populations. Currently, genetic and proteomic approaches are highly reported to detect metabolic disorders that also include inborn error of metabolisms. These existing detection methods are faced with cost issue and time consuming factors. Therefore, metabolites as biomarkers are one of potential avenues to detect metabolic disorders. Further, exploitation of urine as potential source of metabolite biomarkers, there are limitation in this area of research due to abundance of non-metabolite components such as proteins and nucleic acids. Hence, methods and processes are required to precisely fractionate metabolites from urine of inborn error of metabolism patients and then identified by analytical tools such as LC-HRMS and GC-MS.

**Methods:** Sterile filtered urine samples (750 µl) mixed with (250 µl) loading buffer were electrophoresed on VTGE that uses acrylamide gel (acrylamide:bisacrylamide, 30:1) as matrix of 15%. Further, vertical tube gel electrophoresis (VTGE) technique combined with LC-HR-MS to identify metabolites that are known as the biomarkers of metabolic disorders was carried out.

**Results and Discussion:** The authors provide evidence on the use of novel VTGE coupled with LC-HRMS to detect metabolites among metabolic disorders. Data suggest the applicability of VTGE coupled with LC-HRMS technique to detect metabolites such as 2-methyluridine, 2-Methylglutaric acid, 2-Methyl citric acid, 2-Hydroxyglutaric acid in case of metabolic disorders.

**Conclusion:** This preliminary work is suggested to be extended to large clinical samples to validate application of this method to detect metabolic disorders including inborn error of metabolisms.

## INTRODUCTION

Among various classes of metabolic disorders, organic acidurias a type of inherited metabolic disorders is known to surface as intermediate steps in a metabolic pathways of anabolism and catabolism of lipid, carbohydrate, nucleic acids and amino acids (1-5). In essence, these metabolic disorders are indicated to lead to abundance of organic acids including 2-methyluridine, 2-Methylglutaric acid, 2-Methyl citric acid, 2-Hydroxyglutaric acid in a relevant tissues and these metabolites are potentially excreted in urine (5-13).

Among various classes of metabolic disorders, inborn errors of metabolism are seen as one of a kind of genetic disorders. In this class of metabolic disorders, lack or altered enzyme activity can lead to the disrupted biochemical processes that lead to the pathophysiological conditions (5-13). In essence, such metabolic disorders display a condition of either as the aberrant abnormal accumulation of a substrate and on the other hand, a deficit of the product is noticed in a target clinical patient. Further, it is also understood that all such metabolic disorders are inherited in an autosomal recessive manner (5-13). Based on literature, more than 500 human diseases depicted as the metabolic disorders are reported in clinical settings. Predominantly, these inborn errors of metabolism are shown to affect more than one children among 1,000 population. Interestingly, in the Indian settings, lack of awareness and established method to adopt genetic and metabolic approaches, these metabolic disorders are remained unnoticed and undetected. Such unnoticed and undetected metabolic disorders from early stage of life to adult stage are documented to be implicated in many human diseases including diabetes and cancer (10-13). Hence, metabolic screening is recommended during early stage of life and may also be extended to later part of life. Among various markers, metabolic acidosis are suggested as one of important display during such metabolic disorders (10-13). Currently, various modalities including genetic and metabolomics approaches are reported in literature. However, these methods and processes are faced with limitations in terms of validation accuracy and feasibility of translation at bedside approaches.

In this paper, the authors present an approach that combines the vertical tube gel electrophoresis (VTGE) based fractionation of urine samples and liquid chromatography-high resolution mass spectrometry (LC-HRMS) identification of metabolites in the urine samples of potential human subjects on a pilot experimental basis.

## MATERIALS AND METHODS

### Collection and preparation of urine samples

Fresh urine samples from healthy clinical subjects were collected in sterile collection tube. A formal approval was obtained through institutional ethics committee (IEC) and institutional scientific committee (ISC) in accordance with the standard protocol to conduct research on healthy clinical subjects. An informed consent was obtained from the participating healthy clinical subjects. The Further, these urine samples were centrifuged twice at 12000Xg for 30 min. to get rid of debris, dead cells, insoluble particulates and other interfering precipitates. Next, centrifuged urine samples were filtered using 0.45 micron syringe filter to get clear and sterile urine filtrate for subjected to VTGE based fractionation of metabolites.

### VTGE based fractionation of metabolites from urine

Sterile filtered urine samples (750 µl) mixed with (250 µl) loading buffer were electrophoresed on VTGE that uses acrylamide gel (acrylamide:bisacrylamide, 30:1) as matrix of 15%. The fractionated elute was collected in electrophoresis running buffer that contains water and glycine and excludes traditional SDS, and other reducing agents. A flow diagram of VTGE method is presented in Figure 1A and 1B that shows the assembly and design of VTGE system (14-15).

**Figure 1.**
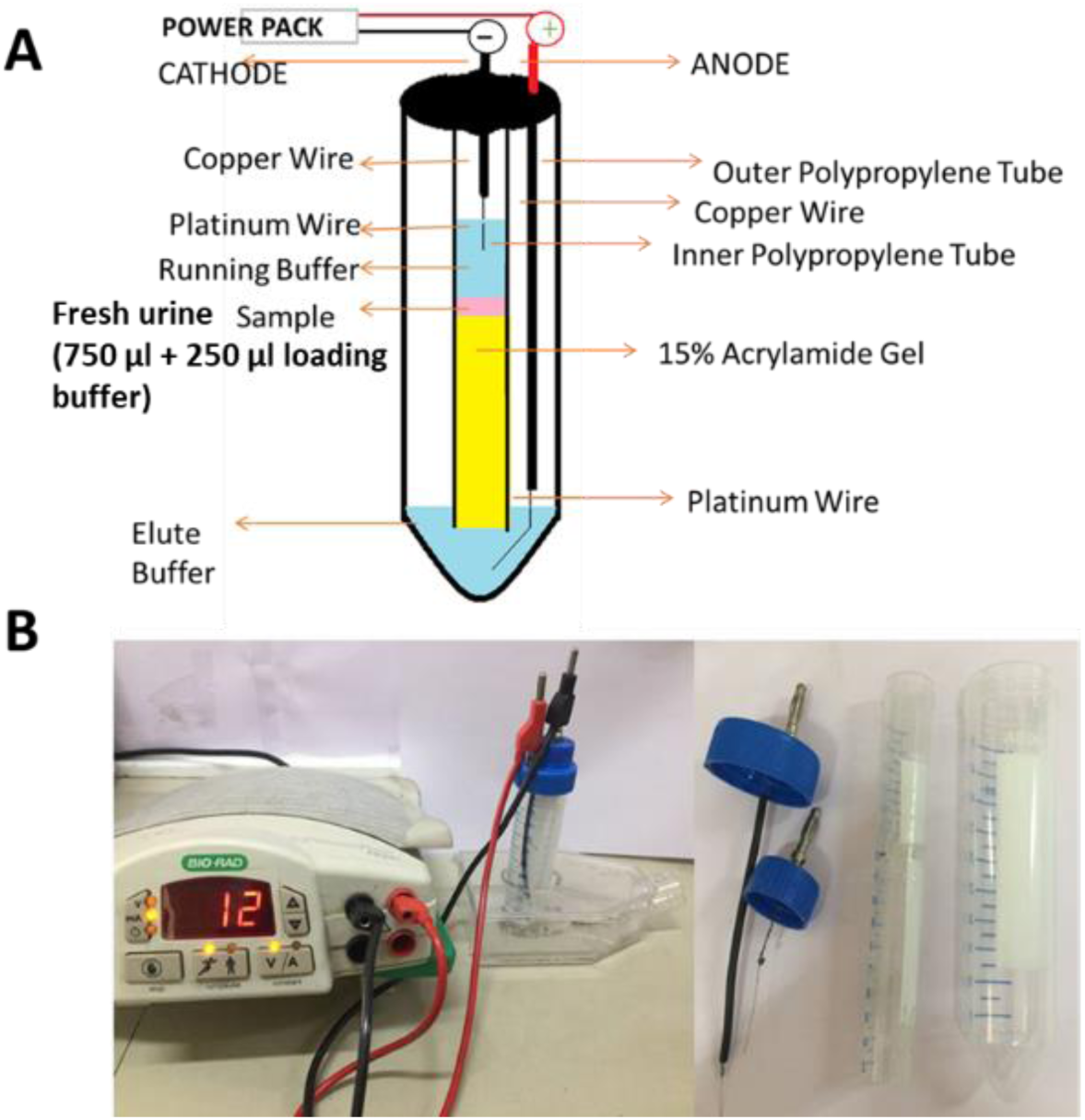
Assembly and working of VTGE system to elute urine metabolites

Urine metabolites from healthy clinical subjects were eluted in the same running buffer, which is referred as elution buffer. After loading of urine samples along with loading buffer, power supply was connected and voltage and current ratio were maintained to generate 1500-2500 mW of power to achieve the electrophoresis of urine samples. The total run time has been allowed for 2 hr and at the end of 2 hr, lower collecting buffer was collected in tube for LC-HRMS characterization. At the end of run, inner tube containing polyacrylamide gel was removed and put for coomassie brilliant blue dye staining to ensure that protein components of urine samples were trapped in the PAGE gel. Further, these eluted fraction from different healthy clinical subjects were stored at −20°C for LC-HRMS analysis and identification of urine metabolites.

### LC-HR-MS analysis of VTGE fractionated urine metabolite elute

LC-HRMS analysis of VTGE metabolite elutes were performed in negative electrospray ionization (ESI) M-H adduct mode. For the analysis of urine metabolites in LC-HRMS, a flow rate of 0.2 mL/min and a gradient was formed by mixing mobile phase A (water containing 5mM ammonium acetate) and B (0.2% formic acid). The HPLC column effluent was allowed to move onto an Electrospray Ionization Triple Quadrupole Mass Spectrometer (Agilent Technologies). Here, urine metabolites were analyzed by using negative electrospray ionization in the multiple reaction monitoring mode (1-3, 15). During LC-HR-MS analysis of urine metabolites, mass spectrometer component was used as MS Q-TOF Quadrupole time-of-flight mass spectrometry (Q-TOF-MS) (Agilent Technologies, 6500 Series Q-TOF LC/MS System) with dual EJS electrospray ionization (ESI) mode. For liquid chromatography (LC) component, RPC18 column as GB metabolite, column Zorbax, 2.1×50 mm, 1.8 micron meter was used to separate the urine metabolite components. The acquisition mode of MS1 is maintained at minimum range of m/z at 60 and maximum range of m/z at 1700. For the run of sample, an injection volume was maintained at 25 µl and a flow rate of solvent was at 0.3 ml per minute.

## RESULTS

In view of need for the detection of potential metabolites of metabolic disorders, a VTGE method combined with LC-HRMS is used to detect these metabolites as biomarkers in urine samples of healthy clinical subjects. In this paper, VTGE method that employs 15% polyacrylamide gel matrix to fractionate in the range of less than 500 Da. metabolites derived from urine samples. These fractionated urine metabolites were subjected to LC-HRMS to detect the nature of metabolites. Here, electrospray ionization chromatogram (EIC) of 3-methyluridine (Figure 2), 2-methylcitric acid (Figure 3) and 2-methyglutaric acid (Figure 4) and 2-hydroxyglutaric acid (Figure 5).

**Figure 2.**
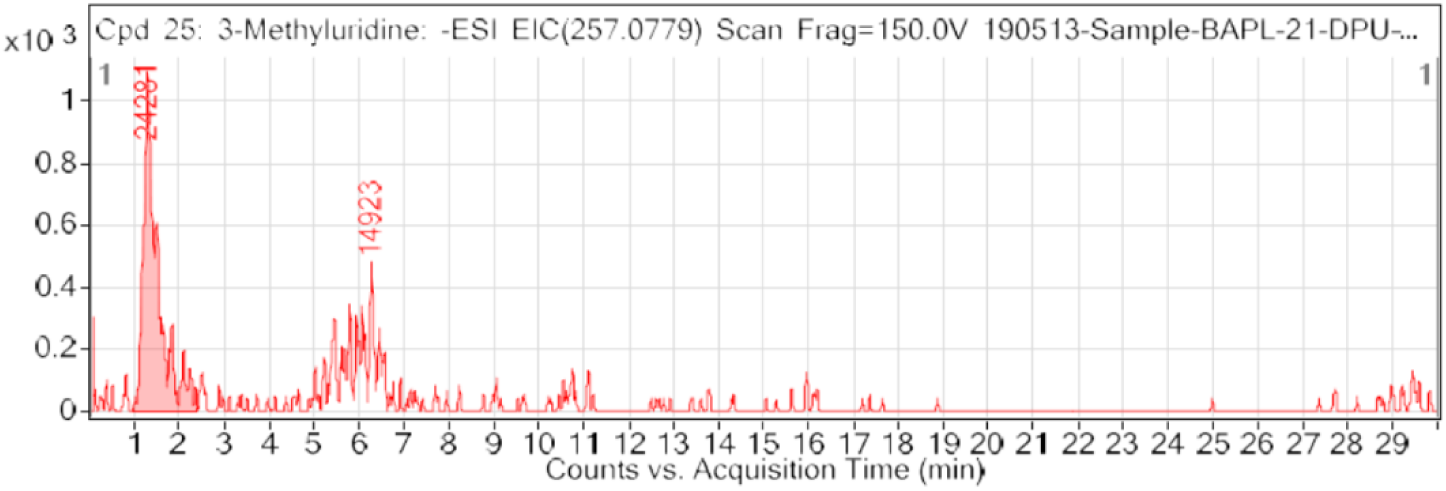
A LC-HRMS based electrospray ionization chromatogram (EIC) of VTGE fractionated urine metabolites from healthy clinical subjects. In this EIC, 3-methyluridine is detected as potential urine metabolite.

**Figure 3.**
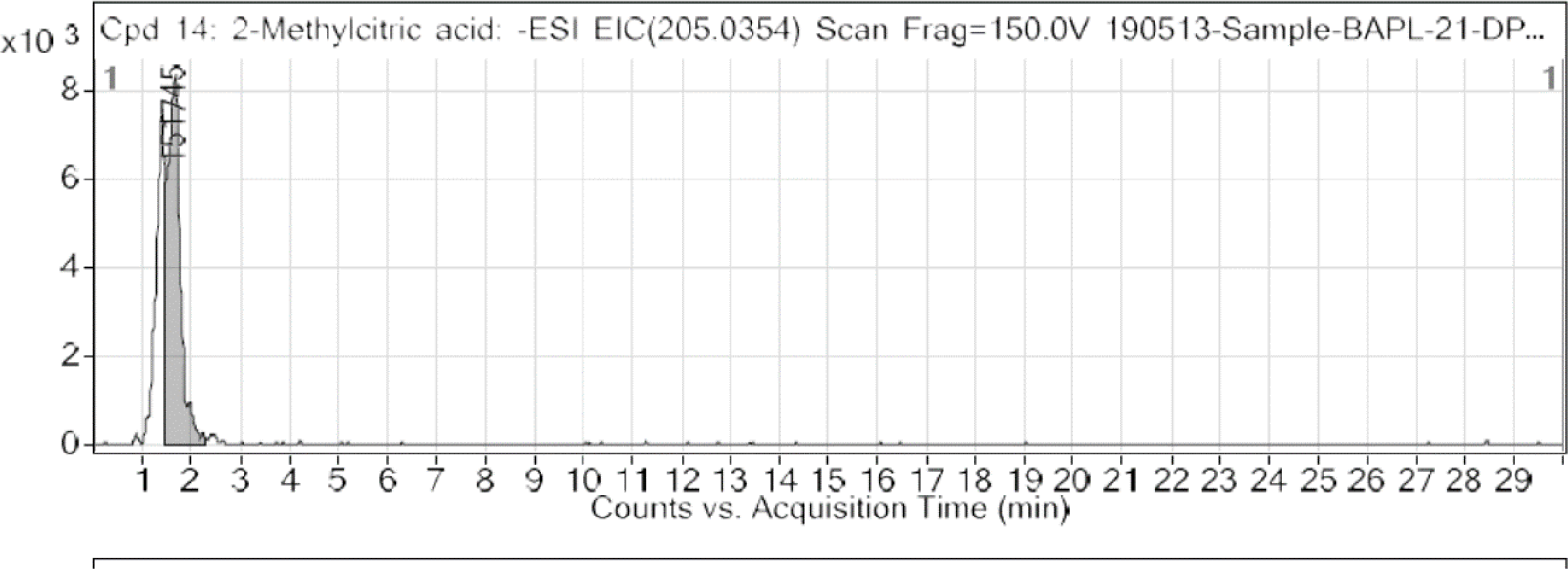
A LC-HRMS based electrospray ionization chromatogram (EIC) of VTGE fractionated urine metabolites from healthy clinical subjects. In this EIC, 2-methylcitric acid is shown as potential urine metabolite.

**Figure 4.**
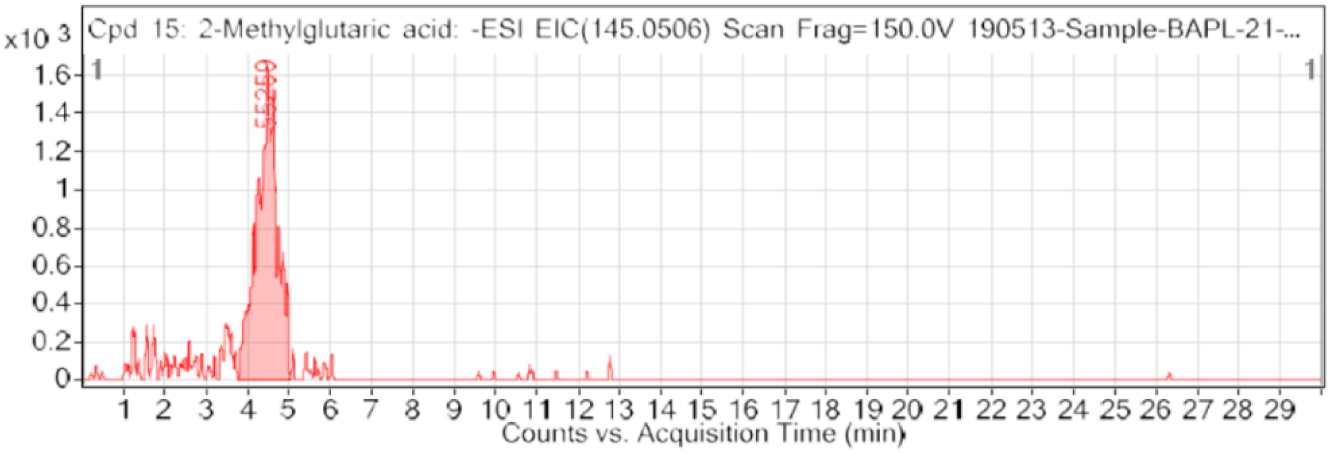
A LC-HRMS based electrospray ionization chromatogram (EIC) of VTGE fractionated urine metabolites from healthy clinical subjects. In this EIC, 2-methyglutaric acid is shown as potential urine metabolite.

**Figure 5.**
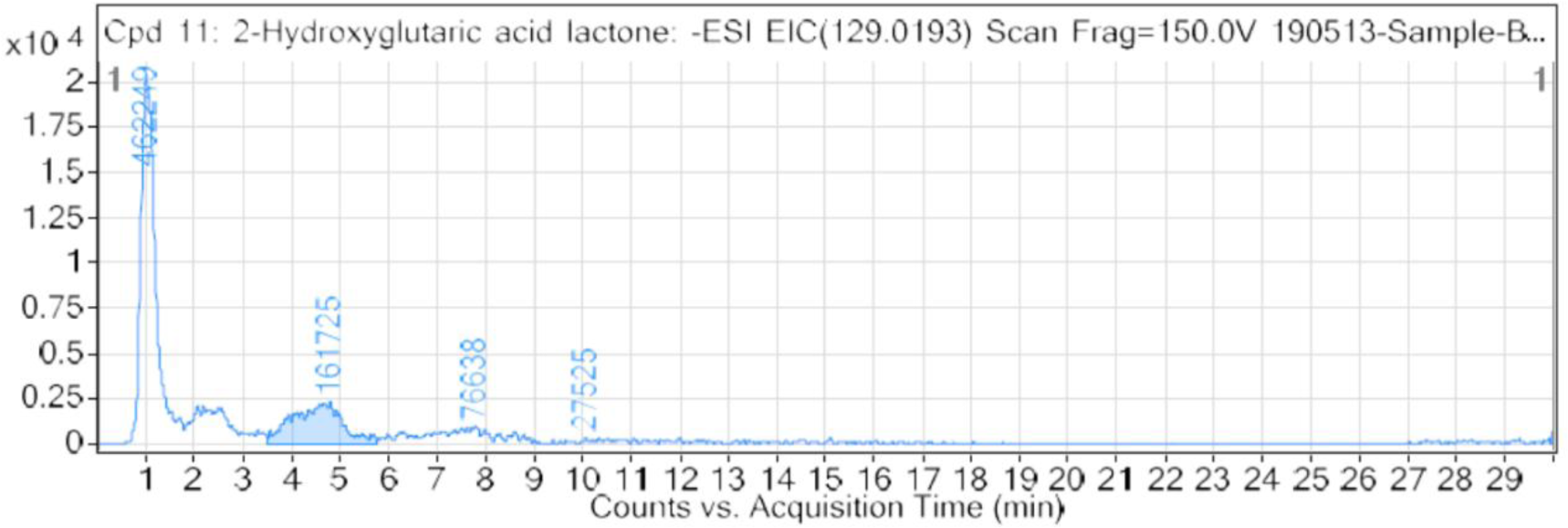
A LC-HRMS based electrospray ionization chromatogram (EIC) of VTGE fractionated urine metabolites from healthy clinical subjects. In this EIC, 2-hydroxyglutaric acid is shown as potential urine metabolite.

A detailed analysis of molecular structure, mass and polarity of these potential metabolites is given in Table 1. This table clearly signifies the detection of 3-methyluridine, 2-methylcitric acid, 2-methyglutaric acid and 2-hydroxyglutaric acid. In this paper, all detected metabolites are in the range of 130-258 Da. M.W. In other way, this data substantiate the working of VTGE method that is configured to fractionate metabolites with less than 500 Da. M.W.

**Table 1:** List of detected urine metabolites as potential biomarkers of metabolic disorders

| Sr No | Name of metabolites | Formula | RT | m/z | Mass | Score | Polarity |
| --- | --- | --- | --- | --- | --- | --- | --- |
| 1. | 2-Methylglutaric acid | C<sub>6</sub> H<sub>10</sub> O<sub>4</sub> | 4.479 | 145.0507 | 146.0571 | 72.22 | Negative |
| 2. | 2-Hydroxyglutaric acid lactone | C<sub>5</sub> H<sub>6</sub> O<sub>4</sub> | 1.003 | 129.0185 | 130.0256 | 83.92 | Negative |
| 3. | 2-Methylcitric acid | C<sub>7</sub> H<sub>10</sub> O<sub>7</sub> | 1.394 | 205.034 | 206.0413 | 90.81 | Negative |
| 4. | 3-Methyluridine | C<sub>10</sub> H<sub>14</sub> N<sub>2</sub> O<sub>6</sub> | 1.286 | 257.0789 | 258.0864 | 72.08 | Negative |

## DISCUSSION

In human metabolome database, 2-methylglutaric acid is known as a derivative of leucine metabolite (1-13). It is suggested that secretion of methylglutaric acid can be found in the urine of patients showing lack of 3-methylglutaconyl coenzyme A hydratase and 3-hydroxy-3-methylglutaryl-CoA lyase deficiency during a type of inborn error of metabolisms. Another example of metabolite as 2-Hydroxyglutaric acid is known to be accumulated in an organic aciduria condition (1-5). There is an indication that 2-hydroxyglutaric acid can be generated due to the enzymatic action of hydroxyacid-oxoacid transhydrogenase and further 2-Hydroxyglutaric acid is shown to be converted into alpha-ketoglutaric acid by 2-hydroxyglutarate dehydrogenase. There is a known fact that 2-Hydroxyglutaric acid is produced due to the gain-of-function mutations of isocitrate dehydrogenase enzyme and this enzyme is a part of TCA cycle (1-7).

It is interesting to note that 2-hydroxyglutaric acid may inhibit a range of enzymes such as histone lysine demethylases (KDMs) and members of the ten-eleven translocation (TET) family of 5-methylcytosine (5mC) hydroxylases (1-5). In summary, the uses of 2-hydroxyglutaric acid as metabolite biomarkers is having relevance in metabolic disorders and also as a key oncometabolites. In fact, 2-hydroxyglutaric acid is known as an alpha hydroxy acid that belongs to the class of organic compounds as organic acids. There are findings that support that levels of 2-hydroxyglutaric acid are an indicator of inborn error of metabolism as an organic aciduria. Organic aciduria can be found in both children and adults that shows various symptoms as tremors, sleepiness, headaches, feeling tired, and seizures (1-13). In line with existing evidence on the potential use of 2-hydroxyglutaric acid, 2-methylglutaric acid, the authors report on a novel methods and processes to detect these urine metabolites as a source of biomarkers for metabolic disorders.

A gas chromatography-tandem mass spectrometry (GC-MS/MS), technology based metabolite biomarkers of inborn error of metabolic disorders that report on the detection of methylmalonic academia (Wang et al., 2017). In the same direction, another LC-MS/MS based detection of methylmalonic acid, 2-methylcitric acid, and homocysteine are reported in case of potential cases of organic aciduria among infant patients (Want et al., 2019). Other than urine sample, a mass spectrometry approach is reported on the detection of methylmalonic acid, 2-methylcitric acid, and homocysteine in a blood spot samples (Turgeon et al., 2010). A quantitative approach to estimate hydroxyglutaric acid in plasma and urine is reported by using LC-MS/MS system to confirm the glutaric aciduria among paients (Simon and Wierenga, 2018). In convergence of earlier reported GC-MS and LC-MS based approaches to identify, our data is promising by adopting a new methods by combining VTGE coupled with LC-HRMS for the detection of urine metabolites as potential biomarkers for metabolic disorders.

## CONCLUSION

This paper reports on the use of VTGE combined with LC-HRMS technique for the identification of 2-methyluridine, 2-Methylglutaric acid, 2-Methyl citric acid, 2-Hydroxyglutaric acid as potential metabolite biomarkers in case of metabolic disorders clinical subjects. This is report is of novel in the sense that uses new method VTGE to fractionate metabolites from urine samples and further identifications of potential metabolites relevant to metabolic disorders. This study is limited to less number of clinical subjects, however, may be extended to large sample size for application in clinics for the detection of metabolic disorders in children and adult subjects.

## GENERATION OF THE INTERDISCIPLINARY WORK

This work is interdisciplinary in nature by collaborating between Department of Biotechnology and Department of Oral Pathology. This reported work is a part of pilot study to characterize urine metabolites relevant to human diseases including cancer. Dr. Nilesh Kumar Sharma conceived the idea of VTGE method and process and established clinical collaboration with Prof. Sachin Sarode, who is a lead investigator from the Department of Pathology.

### Sources of financial support, technical or other form of support if any

DPU funded project to Dr. Nilesh K Sharma and DPU Innovation award (2019) to a team lead by Dr. Nilesh K. Sharma.

### Declaration of conflicts of interest

The authors declare none conflict of interests.

## Acknowledgements

The authors acknowledge inputs and suggestions from Prof. J. K. Pal, Dr. Ramesh Bhonde and assessment team of DPU Innovation award (2019) to improve VTGE system for better applicability in health research.

## References

1. Simon GA, Wierenga A. Quantitation of plasma and urine 3-hydroxyglutaric acid, after separation from 2-hydroxyglutaric acid and other compounds of similar ion transition, by liquid chromatography-tandem mass spectrometry for the confirmation of glutaric aciduria type 1. J Chromatogr B Analyt Technol Biomed Life Sci. 2018. 1097-1098:101–110.

2. Stenton SL, Kremer LS, Kopajtich R, Ludwig C, Prokisch H. The diagnosis of inborn errors of metabolism by an integrative “multi-omics” approach: A perspective encompassing genomics, transcriptomics, and proteomics. J Inherit Metab Dis. 2019. doi: 10.1002/jimd.12130.

3. Shanmuganathan M, Britz-McKibbin P. New Advances for Newborn Screening of Inborn Errors of Metabolism by Capillary Electrophoresis-Mass Spectrometry (CE-MS). Methods Mol Biol. 2019. 1972:139–163.

4. Villani GR, Gallo G, Scolamiero E, Salvatore F, Ruoppolo M. “Classical organic acidurias”: diagnosis and pathogenesis. Clin Exp Med. 2017. 17(3):305–323.

5. Turgeon CT, Magera MJ, Cuthbert CD, Loken PR, Gavrilov DK, Tortorelli S, Raymond KM, Oglesbee D, Rinaldo P, Matern D. Determination of total homocysteine, methylmalonic acid, and 2-methylcitric acid in dried blood spots by tandem mass spectrometry. Clin Chem. 2010. 56(11):1686–95.

6. Wang Y, Sun Y, Jiang T. Clinical Application of LC-MS/MS in the Follow-Up for Treatment of Children with Methylmalonic Aciduria. Adv Ther. 2019. 36(6):1304–1313.

7. Valik D, Jones JD. Hereditary disorders of purine and pyrimidine metabolism: identification of their biochemical phenotypes in the clinical laboratory. Mayo Clin Proc. 1997. 72(8):719–25.

8. Wang H, Wang X, Li Y, Dai W, Jiang D, Zhang X, Cui Y. Screening for inherited metabolic diseases using gas chromatography-tandem mass spectrometry (GC-MS/MS) in Sichuan, China. Biomed Chromatogr. 2017. 1(4).

9. Schillaci LP, DeBrosse SD, McCandless SE. Inborn Errors of Metabolism with Acidosis: Organic Acidemias and Defects of Pyruvate and Ketone Body Metabolism. Pediatr Clin North Am. 2018. 65(2):209–230.

10. Garrod AG. Inborn error of metabolism. Oxford: Oxford University Press; 1909.

11. Scriver CR, Beaud, Sly WS, Valle D, et al., editors. The metabolic and molecular bases of inherited disease. 8th ed. New York: McGraw-Hill; 2001.

12. Superti-Furga A, Hoffman GF. Glutaric aciduria type 1 (Gutaryl CoA-dehydrogenase deficiency): Advances and unanswered questions. Eur J Pediatr. 1997;157:821.

13. Lee HJ, Kremer DM, Sajjakulnukit P, Zhang L, Lyssiotis CA. A large-scale analysis of targeted metabolomics data from heterogeneous biological samples provides insights into metabolite dynamics. Metabolomics. 2019. 15(7):103.

14. Sharma NK. Kumar A, Waghmode A. 2019. Design of vertical tube electrophoretic system and method to fractionate small molecular weight compounds using polyacrylamide gel matrix. Date of Publication: 01/03/2019. (Patent Application Number no: 201921000760). Publication Type INA, The patent official Journal No-19/2018, Page no-9035. Published.

15. Sharma NK, Sarode SC, Pal R. 2019. “A method of urine metabolite profiling by combining vertical tube gel electrophoresis and LC-HR-MS for the detection of oral cancer”. Date of filing 2019/05/29. (Ref. No: 201921021395). Filed/Published.

